# Relative Abundance of SARS-CoV-2 Entry Genes in the Enterocytes of the Lower Gastrointestinal Tract

**DOI:** 10.1101/2020.04.08.033001

**Authors:** Jaewon J. Lee, Scott Kopetz, Eduardo Vilar, John Paul Shen, Ken Chen, Anirban Maitra

**Affiliations:** Department of Translational Molecular Pathology; Sheikh Ahmed Center for Pancreatic Cancer Research; Department of Surgical Oncology; Department of Gastrointestinal Medical Oncology; Department of Clinical Cancer Prevention; Department of Bioinformatics and Computational Biology, The University of Texas MD Anderson Cancer Center

**Author notes:** **Correspondence:** Anirban Maitra, M.B.B.S., Sheikh Ahmed Center for Pancreatic Cancer Research, Department of Translational Molecular Pathology, The University of Texas MD Anderson Cancer Center, 6565 MD Anderson Blvd, Houston, TX 77030.

## Abstract

COVID-19, the disease caused by severe acute respiratory syndrome coronavirus 2 (SARS-CoV-2), has rapidly spread throughout the world and was declared a pandemic by the World Health Organization, thus leading to a rapid surge in the efforts to understand the mechanisms of transmission, methods of prevention, and potential therapies. While COVID-19 frequently manifests as a respiratory infection,^1^ there is evidence for infection of the gastrointestinal (GI) tract^1–4^ with documented viral RNA shedding in the stool of infected patients.^2,4^ In this study, we aimed to investigate the expression of *ACE2* and *TMPRSS2*, which are required for SARS-CoV-2 entry into mammalian cells,^5^ from single-cell RNA sequencing (scRNA-seq) datasets of five different parts of the GI tract: esophagus, stomach, pancreas, small intestine, and colon/rectum.

## Methods

We utilized previously published scRNA-seq data, which are summarized in **Supplementary Table 1**. For in-house normal colon samples, a total of 7 patients were recruited at the University of Texas MD Anderson Cancer Center through written informed consent following Institutional Review Board approval. Single cell dissociation, library generation, and sequencing were performed following standard protocols described in **Supplementary Materials**.

## Results

Analysis of scRNA-seq was performed separately for each GI segment and available cell identities from the original studies were assigned to the new clusters after confirming the expression of relevant marker genes (**Figure 1**). Overall, there were 82,422 cells from the esophagus, 18,405 cells from the stomach, 13,407 cells from the pancreas, 9,858 cells from the small intestine, and 20,161 cells from the colon and rectum (**Supplementary Table 1**). The proportions of cells expressing *ACE2* were approximately 16-fold lower in the upper GI tract (esophagus, stomach; 1% or 1,004/100,827 cells) than in the lower GI tract (ileum, colon, rectum; 16% or 4,777/30,019 cells), and overall, higher proportions of cells expressed *TMPRSS2* than *ACE2* throughout the GI tract (**Supplementary Figures 1A-E**).

**Figure 1.**
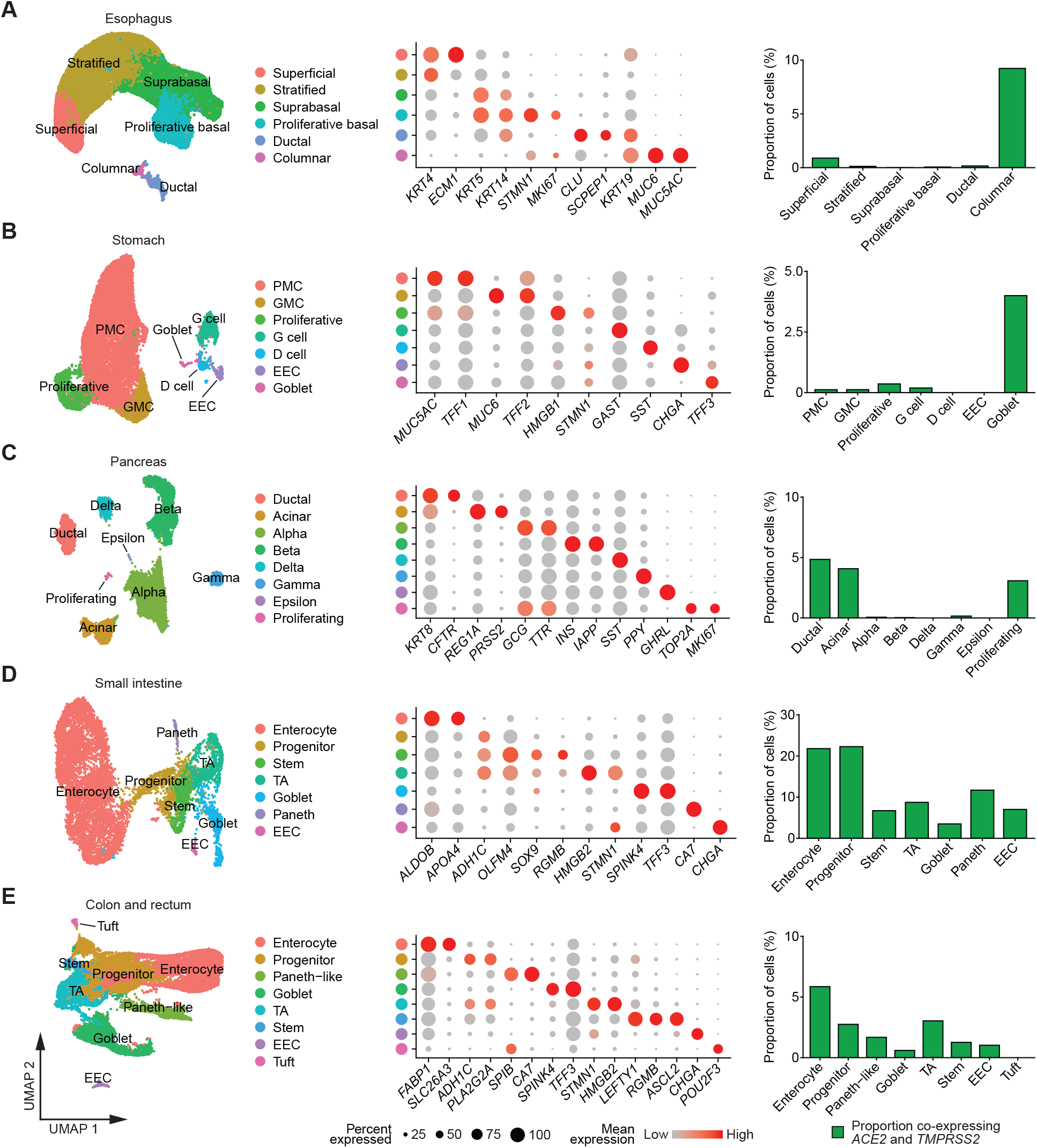
Uniform manifold approximation and projection plots (left) of scRNA-seq data from esophagus (**A**), stomach (**B**), pancreas (**C**), small intestine (**D**), and colon and rectum (**E**); bubble plots showing expression levels of cell-type marker genes (middle); and bar plots depicting proportions of cell types co-expressing *ACE2* and *TMPRSS2* (right). EEC: enteroendocrine cell; GMC: antral basal gland mucous cell; PMC: pit mucous cell; TA: transit-amplifying cell.

Of note, higher proportions of esophageal columnar cells – which were mostly from Barrett’s esophagus samples – co-expressed *ACE2* and *TMPRSS2*, compared to the native squamous epithelium (**Figure 1A**). Pancreatic ductal and acinar cells co-expressed *ACE2* and *TMPRSS2* but endocrine cells did not show detectable co-expression (**Figure 1C**). Parenthetically, the expression of *TMPRSS2* in acinar cells underscores the rationale for using a TMPRSS2 inhibitor (camostat mesylate) in acute pancreatitis. In fact, this agent is now undergoing early phase clinical trials in COVID-19 patients.^5^

Taken together, within the GI tract, the co-expression of *ACE2* and *TMPRSS2* transcripts was highest in the small intestine and colon. Approximately 20% of enterocytes from the small intestine and 5% of colonocytes (enterocytes from colon/rectum) co-expressed *ACE2* and *TMPRSS2* (**Figure 1D–E**). Analysis of 308 genes that were positively correlated (Pearson’s *r* > 0.1) with *ACE2* in the small intestine revealed the enrichment of functional Gene Ontology (GO) terms such as metabolic and transport pathways in the small intestine (**Supplementary Figure 1F**), thus confirming the inherent digestive and absorptive functions of enterocytes. Interestingly, 251 genes positively correlated with *ACE2* in the colon and rectum showed enrichment of not only secretion-associated pathways but also immune-related processes (**Supplementary Figure 1G**), which may be upregulated in functional colonocytes due to the microbiome.

## Discussion

To summarize, our results confirm the co-expression of SARS-CoV-2 entry genes *ACE2* and *TMPRSS2* in a subset of epithelial cells in the GI tract, especially functional enterocytes from the lower GI tract (small intestine and colon), and provide potential insights into why entero-colitic symptoms may arise in acute COVID-19 infection. These findings parallel the findings from recent studies that revealed the co-expression of *ACE2* and *TMPRSS2* in olfactory epithelium^6^ and intrahepatic cholangiocytes,^7^ which may explain atypical COVID-19 symptoms and laboratory findings such as anosmia/ageusia and transaminitis. Our results further support the feasibility of SARS-CoV-2 entry into the GI tract with implications for GI infection and transmission.

## Acknowledgements

We thank Charles Bowen for performing single cell dissociations from colonic samples, Kangyu Lin and Jinzhuang Dou for assistance in data acquisition and transfer, and the Single Cell Core at MD Anderson Cancer Center.

## References

1. Guan WJ, Ni ZY, Hu Y, et al. N Engl J Med 2020 [E-pub ahead of print].

2. Holshue ML, et al. N Engl J Med 2020;382:929–936.

3. Xiao F, Tang M, Zheng X, et al. Gastroenterology 2020 [E-pub ahead of print].

4. Wolfel R, Corman VM, Guggemos W, et al. Nature 2020 [E-pub ahead of print].

5. Hoffmann M, Kleine-Weber H, et al. Cell 2020 [E-pub ahead of print].

6. Brann DH, Tsukahara T, Weinreb C, et al. bioRxiv 2020.

7. Wen Seow JJ, et al. bioRxiv 2020.

