## Supplementary Materials for "Relative Abundance of SARS-CoV-2 Entry Genes in the Enterocytes of the Lower Gastrointestinal Tract"

### Supplementary Methods

Publicly available single-cell sequencing datasets from esophagus,[1,2] stomach,[3] pancreatic islets,[4-7] small intestine,[8,9] and colon/rectum[10] specimens were obtained as described in **Supplementary Table 1**. For in-house data, single cell dissociation was performed following a standard protocol: tissues were minced with sterile surgical scalpels to approximately 1-mm fragments and resuspended in 0.5 mg/ml Liberase TH (Sigma) followed by incubation at 37°C for 15 min with constant agitation. Liberase was quenched with equal volume of 1% BSA and cells were resuspended in Accutase (Sigma) followed by incubation at 37°C for 15 min with constant agitation. Dissociated cells were passed through a 40 μm strainer and resuspended in 0.04% BSA for subsequent viability analysis and counting. Library generation and sequencing were performed using the 3' Library Construction Kit (10x Genomics) following the manufacturer's recommendations. Single cell data processing was done using standard Cell Ranger RNA pipeline (10x Genomics) using hg19 as a reference.

All downstream single cell analyses were performed in R v3.6.2 using Seurat v3.1.0.[11] Preprocessing included removal of genes expressed in less than 3 cells, removal of cells containing less than 200 genes, and removing cells with high percentage of mitochondrial genes based on distribution with general cutoff ranging 20-50%. Given the size of the datasets, integration was performed using reciprocal principal component analysis as a part of standard Seurat workflow.[11] Log-count greater than zero was defined as presence of expression. Gene Ontology (GO) enrichment analysis was done on geneontology.org[12,13] using genes that were positively correlated with *ACE2* in all small intestine and colorectal cells with Pearson's $r > 0.1$, which were obtained using cor() function.

**References**


1. Owen RP, White MJ, Severson T, et al. Nat Commun 2018;9:4261.
2. Madissoon E, Wilbrey-Clark A, et al. Genome Biol 2019;21:1.
3. Zhang P, et al. Cell Rep 2019;27:1934-1947 e5.
4. Baron M, Veres A, Wolock SL, et al. Cell Syst 2016;3:346-360 e4.
5. Muraro MJ, Dharmadhikari G, et al. Cell Syst 2016;3:385-394 e3.
6. Segerstolpe A, Palasantza A, et al. Cell Metab 2016;24:593-607.
7. Xin Y, et al. Cell Metab 2016;24:608-615.
8. Wang Y, Song W, et al. J Exp Med 2020;217.
9. Martin JC, et al. Cell 2019;178:1493-1508 e20.
10. Parikh K, Antanaviciute A, Fawkner-Cobett D, et al. Nature 2019;567:49-55.
11. Stuart T, Butler A, et al. Cell 2019;177:1888-1902 e21.
12. Ashburner M, et al. Nat Genet 2000;25:25-9.
13. Mi H, et al. Nucleic Acids Res 2019;47:D419-D426.

**Supplementary Table 1.** Summary of datasets used in the study

| Tissue | Dataset | Number of Cells | Source | Notes |
|---|---|---|---|---|
| Esophagus | Madissoon | 81,176 | Seurat object from Human Cell Atlas | |
| | Owen | 1,246 | Counts matrix from supplementary data | Includes Barrett's esophagus samples |
| | All | 82,422 | | |
| Stomach | Owen | 213 | Counts matrix from supplementary data | |
| | Zhang | 18,192 | Counts matrix (GSE134520) | Includes non-atrophic gastritis and chronic atrophic gastritis |
| | All | 18,405 | | |
| Pancreatic islets | Baron | 7,744 | Single Cell Experiment object from Hemberg Group | All samples were enriched for endocrine cells |
| | Muraro | 2,002 | | |
| | Segerstolpe | 2,121 | | |
| | Xin | 1,540 | | |
| | All | 13,407 | | |
| Small intestine | Martin | 4,573 | Counts matrices (GSE134809) | Excluded diseased samples (Crohn's) |
| | Owen | 155 | Counts matrix from supplementary data | |
| | Wang | 5,130 | Counts matrix (GSE125970) | |
| | All | 9,858 | | |
| Colon/rectum | Parikh | 7,640 | Counts matrices (GSE116222) | Excluded diseased samples (ulcerative colitis) |
| | Wang | 7,501 | Counts matrix (GSE125970) | |
| | MDACC | 5,020 | In-house data | |
| | All | 20,161 | | |

### Supplementary Figure Legends

**Supplementary Figure 1.** Bar plots depicting proportions of cell types in esophagus (**A**), stomach (**B**), pancreas (**C**), small intestine (**D**), and colon/rectum (**E**) expressing *ACE2* (dark blue, top) or *TMPRSS2* (yellow, bottom) and bar plots of top 5 GO terms enriched in genes that are positively correlated with *ACE2* expression in small intestine (**F**) and colon/rectum (**G**). EEC: enteroendocrine cell; FDR: false discovery rate; GMC: antral basal gland mucous cell; PMC: pit mucous cell; TA: transit-amplifying cell.

### Supplementary Figure 1

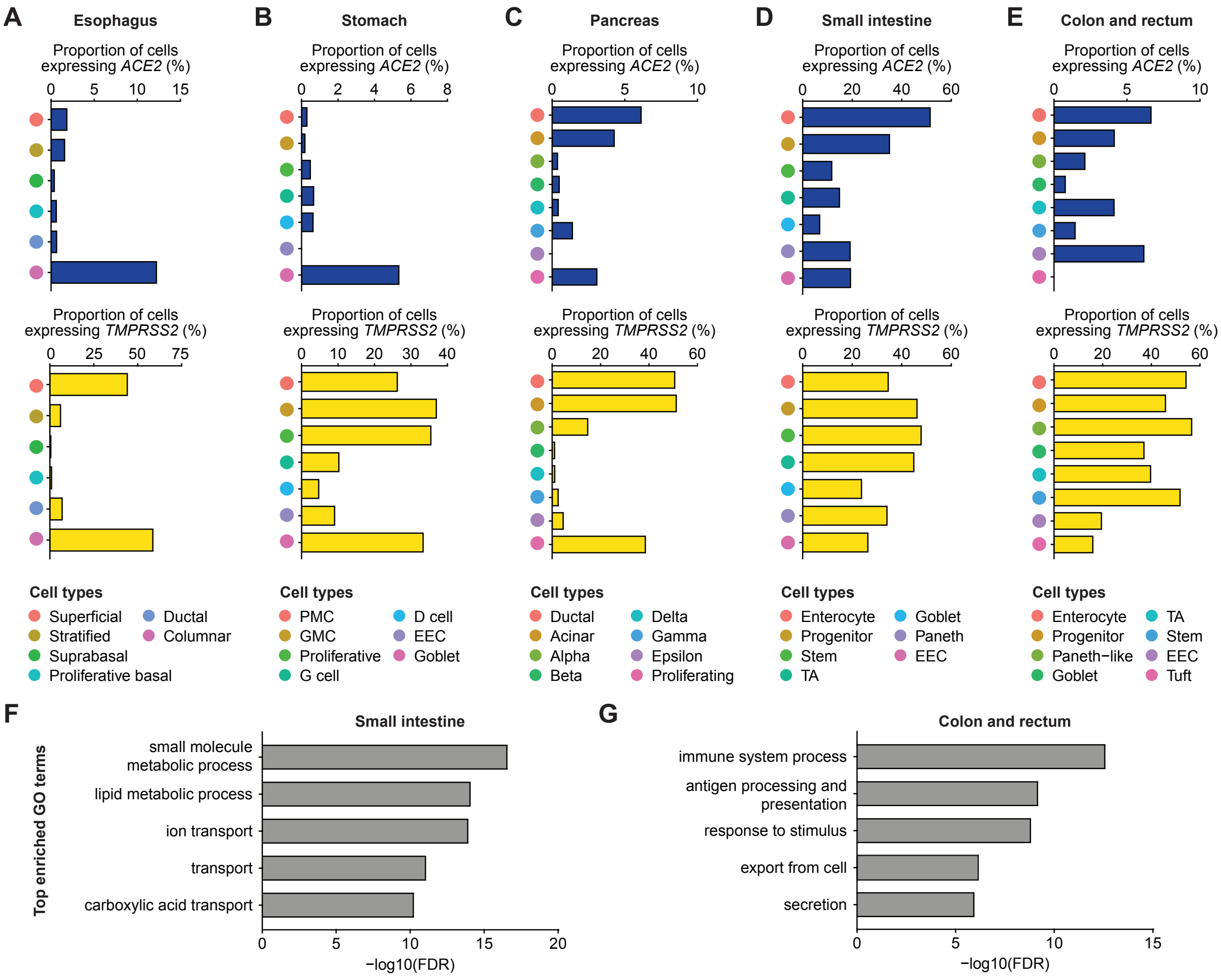
